# A comprehensive water buffalo pangenome reveals extensive structural variation linked to population specific signatures of selection

**DOI:** 10.1101/2025.05.04.652079

**Authors:** Fazeela Arshad, Siddharth Jayaraman, Andrea Talenti, Rachel Owen, Muhammad Mohsin, Shahid Mansoor, Muhammad Asif, James Prendergast

## Abstract

Water buffalo is a cornerstone livestock species in many low- and middle-income countries, yet major gaps persist in its genomic characterization—complicated by the divergent karyotypes of its two sub-species (swamp and river). Such genomic complexity makes water buffalo a particularly good candidate for the use of graph genomics, which can capture variation missed by linear reference approaches. However, the utility of this approach to improve water buffalo has been largely unexplored.

We present a comprehensive pangenome that integrates four newly generated, highly contiguous assemblies of Pakistani river buffalo with available assemblies from both sub- species. This doubles the number of accessible high-quality river buffalo genomes and provides the most contiguous assemblies for the sub-species to date. Using the pangenome to assay variation across 711 global samples, we uncovered extensive genomic diversity, including thousands of large structural variants absent from the reference genome, spanning over 140 Mb of additional sequence. We demonstrate the utility of these data by identifying putative functional indels and structural variants linked to selective sweeps in key genes involved in productivity and immune response across 26 populations.

This study represents one of the first successful applications of graph genomics in water buffalo and offers valuable insights into how integrating assemblies can transform analyses of water buffalo and other species with complex evolutionary histories. We anticipate that these assemblies, and the pangenome and putative functional structural variants we have released, will accelerate efforts to unlock water buffalo’s genetic potential, improving productivity and resilience in this economically important species.

## Background

Water buffalo (*Bubalus bubalis*) are central to the livelihoods of millions of people worldwide, especially in low- and middle-income countries (LMICs). Among these, Pakistan, India, China, and several Southeast Asian nations rely heavily on water buffalo for milk, meat, and draught power. In Pakistan alone, buffaloes contribute around 60% of the total milk production [1]— underscoring their critical role in national food security and the incomes of smallholder farmers. Beyond its contributions to nutrition and the agricultural economy, water buffalo hold cultural importance in many regions, where they represent a valuable, multi-purpose asset that can thrive in diverse ecological conditions.

Despite its socioeconomic significance, water buffalo have historically received less systematic genomic research compared to its bovine relative, the domestic cow (*Bos taurus/Bos indicus*). This is partially due to the lower use of water buffalo in high-income countries, resulting in more limited investment into genomic research and breeding programs. Consequently, key questions remain unanswered, including how different alleles and structural variants (SVs) influence key traits like milk yield, carcass quality, growth rate, and disease resistance. Addressing this could help unlock the genetic potential of water buffalo and dramatically enhance production efficiency, particularly benefiting smallholder farming communities in LMICs.

A major factor complicating water buffalo genomic analyses is the species’ complex evolutionary history. Present-day water buffalo descend from wild *Bubalus arnee*, domesticated about 6,300 years ago in mainland Southeast Asia [2]. Two primary sub-species emerged: river buffalo (2n karyotype=50), widely used for high-yield milk production, and swamp buffalo (2n=48), primarily used for draught and meat [3]. Although they exhibit distinct karyotypes, both sub-species can interbreed and produce fertile offspring [4]. Yet this divergent chromosome number, estimated to have originated around three million years ago, complicates analyses of genomic variation [5]. For example, the current water buffalo reference genome is derived exclusively from a river buffalo [6], which can lead to inaccuracies when aligning swamp buffalo sequences. Swamp populations—that predominate in mainland Southeast Asia—show higher levels of divergence and harbour structural variants not captured by the river-centric reference [5], resulting in biases in variant calling and downstream analyses.

Capturing these large-scale genomic differences is crucial for understanding phenotype variation, such as in relation to milk production, meat quality, and disease resistance. Structural variants (such as insertions, deletions, and inversions) encompass more nucleotides than single nucleotide polymorphisms (SNPs) [7], consequently potentially having larger impacts on heritable phenotypes. Studies in other livestock species have highlight the importance of identifying such variants [8]. However, for water buffalo, the scarcity of high-quality genome assemblies, especially from regions with richly diverse indigenous breeds, has slowed progress. Although eight higher quality long-read-based assemblies are publicly available—[5, 6, 9–12] split evenly between the swamp and river types—none were generated from Pakistani buffaloes, despite the importance of local breeds like Nili-Ravi and Azikheli to milk production in South Asia [13, 14].

Over the past decade, several studies have investigated possible selective sweeps in water buffalo breeds by analysing single nucleotide polymorphisms (SNPs) derived from genotyping arrays or short-read whole-genome sequencing data aligned to a single river buffalo reference [15–18]. These efforts have identified loci putatively linked to economically and biologically important traits, including lactation, fertility, growth, and immune response. For example, Dutta et al. used whole-genome sequencing and population genomic analyses to detect selective signatures in Indian river buffalo, while Sun et al. and Si et al. similarly reported candidate genomic regions under positive selection in both swamp and river lineages. Such studies highlight how domestication pressures and local adaptations have shaped the water buffalo genome in different geographical contexts. Despite these advances, these investigations of selective sweeps in water buffalo have focused on short variation. Whether any larger variants may underlie selective sweeps in the species largely remains unexplored.

Graph genomics offers a powerful alternative to traditional linear reference-based analyses. By integrating multiple assemblies into a single “pangenome,” graph methods can more accurately represent breed- or sub-species–specific haplotypes, including large structural variants [19]. This avoids biases inherent in mapping reads to a single reference, particularly when working with genetically diverse or structurally distinct populations. Graph-based approaches have reduced false-negative rates in larger variant calling for other livestock [20–22] revealing alleles that were previously hidden when aligned to a single linear reference. Yet, such methods have largely not been applied to water buffalo, leaving potentially important functional variation in the species uncharted.

Here, we address this gap by generating new, highly contiguous assemblies for Pakistani river buffalo, significantly expanding existing genomic resources. We construct the first comprehensive pangenome that includes these new assemblies alongside publicly available river and swamp buffalo genomes. By genotyping global water buffalo samples against this pangenome, we generate the largest combined water buffalo reference set of structural and single-nucleotide variation to date, and identify previously unidentified structural variants potentially underlying natural and artificial selective sweeps. Collectively, this work has the potential to inform a diverse range of studies, including breeding programs, conservation strategies, and future genomic analyses, ultimately improving water buffalo productivity and resilience worldwide.

## Methodology

### Sample collection and DNA extraction

The animal handling and sample collection protocol was reviewed and approved by the Research Ethics Committee of the National Institute for Biotechnology and Genetic Engineering (NIBGE), Faisalabad, Pakistan, on May 29, 2024. One female Nili Ravi Buffalo from the Punjab province of Pakistan and one Azikheli female buffalo from Swat, a district of the province of Khyber Pakhtunkhwa, Pakistan was selected for genome sequencing (Supplementary Fig. S1, S2). Blood sample collection was conducted under the supervision of trained animal care specialists to minimize stress and ensure the welfare of the animals. Fresh blood was collected from the jugular vein of animals in EDTA-coated tubes and kept cool on ice gel packs for transportation to the laboratory at NIBGE. Genomic DNA was isolated using the Thermo Scientific GeneJET Whole Blood Genomic DNA Purification Mini Kit following the manufacturer’s protocol. Prior to extraction blood was pre-processed by freshly prepared digestion solution-A (1M MgCl2,1M Tris- HCl (pH 7.5),2M Sucrose and Triton X-100) with the aim of increasing the yield. In total 750 µL of thoroughly homogenized blood was transferred to a sterile 1.5 mL microcentrifuge tube, and an equal volume of “Solution- A" was added. The mixture was vortexed and incubated at room temperature for 10 minutes. It was then centrifuged at 11,000 rpm for 45 seconds. After centrifugation, the supernatant was discarded, and the pellet was resuspended in 400 µL of Solution-A. This process of incubation at room temperature for 10 minutes, followed by centrifugation at the same speed for 45 seconds, was repeated until the supernatant became clear. Following the final removal of the supernatant, the pellet was dissolved in 190 µL of PBS to achieve a final volume of 200 µL. This 200 µL WBC suspension in PBS was subsequently used for DNA extraction following the kit protocol, instead of using whole blood.

### Sequencing and Genome Assembly

Extracted DNA was shipped to the Edinburgh Genomics sequencing facility at The University of Edinburgh in the UK and PacBio HiFi sequencing data was generated using PacBio Revio SMRTbell library preparation.

The resulting HiFi data with a quality of Q ≥ 20 were checked with FASTQC (v0.12.1) [23]. A total of 98.2Gbp of sequence data, with a median read length of 15.8kb, was generated for the Azikheli buffalo (AZ0004), representing a mean depth coverage of ∼34x under an assumed genome size of 2.9Gbp. Similarly, 102.5Gbp of sequence data, with a median read length of 17.1kb, was generated for the Nili Ravi buffalo (NR0003), achieving a mean depth coverage of ∼35x. The long HiFi fastq reads were denovo assembled using HiFiasm (v0.24.0-r702) [24] with the -z20 option added to trim 20bp from both ends of the reads, to produce dual contig level assemblies, i.e. two assemblies per animal.

### Construction of pangenome graph

To make a water buffalo pangenome incorporating both subspecies, eight publicly available water buffalo genome assemblies with the best assembly statistics were obtained. The river buffalo assemblies were: UOA_WB_1 [6], NDDB_DH_1, NDDB_SH_1 [10] and CUSA_RVB [11]. The swamp buffalo assemblies were BBCv1.0 [9], PCC_UOA_SB_1v2 [5], CUSA_SWP [11], and Wang_2023 [12]. These genomes were accessed and downloaded from the National Centre for Biotechnology Information (NCBI), China National Gene Bank (CNGB) and National Genomics Data Centre (NGDC) databases. Further information relevant to the accession numbers can be found in the Data Availability Section.

To eliminate potential biases arising from the differences in tools and versions, and to ensure fair comparison between our novel genome assemblies and publicly available genome assemblies, we recalculated the assembly statistics for all genomes using gfastats (v1.3.6) [25].

The PanGenie Snakemake pipeline [26], was used for the construction of the graph genome from our four newly generated haplotype-resolved assemblies AZ0004_hap1 (Azikheli 1), AZ0004_hap2 (Azikheli 2), NR0003_hap1 (Nili Ravi 1), NR0003_hap2 (Nili Ravi 2) and the seven publicly accessed genomes NDDB_DH_1 (Indian Murrah 1), NDDB_SH_1 (Indian Murrah 2), CUSA_RVB (Chinese Murrah), CUSA_SWP (Zhuang female), BBCv1.0 (Chinese swamp),

PCC_UOA_SB_1v2 (Philippines swamp) and Wang_2023 (Zhuang male), to produce a phased pangenome graph represented as a multiallelic, multi-sample VCF file. This process involved the alignment of water buffalo genome assemblies to the reference genome UOA_WB_1 (Mediterranean 2018) with minimap2 [27]. Since the two buffalo sub-species have different chromosome numbers (river 2n=50 and swamp 2n=48) originating from a fusion of river buffalo chromosomes 4 and 9 [28], chromosome 1 of the swamp assemblies were split at the point of the fusion prior to running the PanGenie Snakemake pipeline. The fusion point was identified using PAF alignments, which provided start and end coordinates of the aligned regions. The midpoint of the gap between these alignment regions was calculated to determine the fusion point. The same approach was applied to all swamp assemblies and the Snakemake workflow was then executed with the updated data.

The resultant, multiallelic, multisample variant call file (VCF) was normalized using the bcftools (v1.20) [29] norm function. To quantify the variation specific to each combination of assemblies a support vector (SUPP_VEC) field was introduced into to the VCF file to record the presence (’1’) or absence (’0’) of the alternative allele for each variant in every sample. Bcftools v1.13 [29] was used to query support vector values for variants with lengths over 50bp, ensuring structural variant information was captured. The resulting data were plotted using the UpSetR package [30] to illustrate the size distribution of structural variants across samples. Following this, to calculate the extra sequence lengths in the buffalo pangenome, the normalized graph filtered VCF file was processed to include length annotations by utilizing the vcflength tool within the vcflib (v1.0.12) [31] environment. The insertion lengths for heterozygous or alternative homozygous variants were extracted for each sample of interest by querying the relevant data with bcftools (v1.13). The total lengths contributed by different assemblies were calculated with the help of a Python script.

For the phylogenetic relationship of the assemblies, we generated a mash distances (distance matrix) based phylogenetic tree from the fasta sequences of studied water buffalo assemblies with the cattle reference *Bos* taurus ARS UCD 2.0 [9] genome added as an outgroup, using mashtree (v1.4.6) [32] with the options --reps 100 and --min-depth 0 to increase bootstraps and ignore low abundance K-mers. Tree visualization was done using figtree (v1.4.4) [33].

### Whole genome sequence data across buffalo populations

The structural variants identified in our pangenome graph were genotyped across a larger cohort of 711 whole-genome sequences from diverse global buffalo populations. Whole genome sequences were accessed from projects PRJNA633724 [34], PRJEB39591 [16], PRJNA547460 [18], PRJCA001294[11], PRJNA350833 [35], PRJNA1135737, PRJNA1057008 and PRJNA633919 [17]. This data was downloaded from the NCBI, NGDC and ENA (European Nucleotide Archive) databases. The 711 water buffaloes from 16 countries included 337 domesticated river buffaloes and 374 domesticated swamp buffaloes (Supplementary Table S1). Both subspecies were further classified into subgroups based on their geographical distribution. The river buffaloes were divided into six geographical groups South Asia (SA), Italy (ITA), West Asia (WA), Egypt (EGY), Nepal (SA-NP) and South Bangladesh (BGD-S). Similarly the swamp buffalo populations were categorized into Central China (CHN-CE), Southwestern China (CHN-SW), Northeast Bangladesh (BGD-NE), Southeast China (CHN-SE), Southeast Asia (SEA) and Indonesia (IND).

### PanGenie Genotyping and Data Filtering

To mitigate against poor quality genotyping, we included only samples with Illumina, paired-end mean read coverage greater than 10X resulting in the above-mentioned sample size of 711 individuals. Fastq reads from the NCBI, ENA and NGDC databases were downloaded using enaBrowserTools (v1.7.1) [36]. The variant call files (VCF) for each unique biosamples were generated using PanGenie (v3.0.0) [26]. PanGenie re-genotyped variants provided in the input graph VCF file derived from the 12 assemblies and referenced to the UOA_WB_1 reference assembly, ensuring that the output VCF contains the same variant records as the input, but with genotypes assigned for the sample on which PanGenie is run. The quality of each genotyped VCF was checked by calculating statistical parameters using RTG Tools (v3.12.1) [37]. Resultant genotyped files for each biosample were merged using the ‘--region’ option in bcftools (v1.13). For downstream analyses, variants with >20% missing genotypes and minor allele frequency <5% were excluded using bcftools options “view -i F_MISSING<0.2”, and “view -i ’MAF>0.05”.

### Relatedness

To assess the population structure of the study cohort, the 711 samples were subjected to kinship analysis to exclude related individuals, monozygotic duplicates and sequencing artefacts thereby ensuring an accurate representation of the genetic pool. This was accomplished by measuring kinship coefficients using King 2.1.2 [32] with options --kinship and --degree 3. To identify and filter related individuals, pairwise kinship coefficients were evaluated using a threshold of 0.0625. Individuals with kinship values above this threshold were grouped into connected components, representing clusters of related individuals. Within these clusters, individuals with higher mean read depth coverage were prioritized and categorized in the "keep" group, while those with comparatively lower coverage were placed in the "remove" group and were excluded. This process generated a list of 403 unrelated individuals. Since after the exclusion of certain samples, the MAF can change, unrelated samples were refiltered for MAF>0.05. Before performing PCA and admixture analyses, LD-based pruning was performed using plink 1.9 [38] with the --indep pairwise 50 10 0.2 options with an additional option of --mind 0.20 to exclude samples with a missing genotype rate >0.20. Then we estimated eigenval and eigenvec values using plink 1.9’s --pca option. Admixture (v1.3.0) [39] was run with the number of assumed ancestral populations (K) ranging from 2 to 9, with K = 6 identified as the best model based on cross-validation (CV) values (Supplementary Fig. S3).

### Genotype concordance

To compare variant calls between PanGenie and GATK we used the 81 samples from a previous study for which GATK calls were also available [16]. Note that two samples were technical replicates in this dataset, meaning it contained 79 distinct animals. Variants were hard filtered as previously described. Importantly this did not involve the use of GATK’s VQSR which may bias the results due to its dependence on existing sets of known variants. Genotypes were compared using RTG tools (v 3.12.1) [37] having first applied a further GQ filter of >= 20 to both VCF files. To determine which variants fell within repetitive regions, repeat masker annotation was downloaded from the UCSC website. Hardy-Weinberg equilibrium values and allele frequencies were calculated using bcftools (version 1.19). Allele frequencies of PanGenie and GATK specific variants was calculated only at variants where at least 50 samples had a genotype call in the respective callset and this analysis was restricted to three representative chromosomes (2, 12 and 22).

### Selective sweep analyses

For the selective sweep analyses the 403 samples filtered for relatedness were grouped based on their breed labels, sampling location and position on the PC1 vs PC2 PCA. This resulted in 26 groups with at least six samples, encompassing a total of 282 samples, that were taken forward for analysis (Supplementary Table S2). Following filtering out genotypes with a genotype quality less than 25, the dataset was further restricted to biallelic variants with genotypes present for at least 75% of samples. The resulting VCF was then phased using Beagle (version 5.2 21Apr21.304), and the integrated haplotype score (iHS) and number of segregating sites by length (nSL) statistics calculated for each variant and group using HapBin (v1.3.0) [40] and Selscan (v2.0.3) [41] respectively. Putative sites of selective sweeps were called as previously [16]. For both the iHS and nSL results, scores were first averaged across 100 variant windows and peaks were then called where the absolute of these mean values rose above 1.5 and fell back below 0.5. Genes overlapping these peaks were then identified using a custom R script and the NCBI gene annotations for water buffalo (Bubalus_bubalis-GCA_003121395.1-2020_06-genes.gff3).

The variant allele frequency differences for 26 selected population groups were calculated by first normalizing the variants using bcftools (v1.20) [29]--norm function, annotating them with SnpEff v5.2f (2025-02-07) [42], and adding fill-tags using bcftools (v1.20) with the focus on high- and moderate-impact variants. Selection regions were defined by merging peak coordinates into unique BED files. Within these regions, alternate allele frequencies (aAF) were queried for each population using bcftools (v1.20), highlighting key high- and moderate-impact variants associated with population-specific genetic adaptations.

## Results

### Novel assemblies for Pakistani breeds

To address the current lack of high-quality assemblies for Pakistani water buffalo breeds, we first PacBio HiFi sequenced one female Nili Ravi (ID:NR0003) and one female Azikheli buffalo (ID:AZ0004), two of the most common breeds used in the country [13]. In total 102.5 Gbp (∼35x coverage) and 98.2 Gbp (∼34x) of data respectively was generated. Using the Hifiasm (v0.24.0- r702) assembler we produced a pair of dual assemblies for each animal with contig N50s ranging from 61 to 84Mb. Comparison of the statistics of these novel assemblies to the best currently publicly available genomes confirms that they are among the most contiguous water buffalo genomes generated to date (Table 1, Figure 1A), and the most complete river buffalo genomes, substantially exceeding the current reference in both total size and contiguity. Phylogenetic analysis of these novel and publicly available assemblies confirms the split between swamp and river buffalo, with the new Pakistani assemblies clustering as a group and most closely to the existing Indian river buffalo assemblies, matching their geographic proximity (Figure 1B).

**Figure 1.**
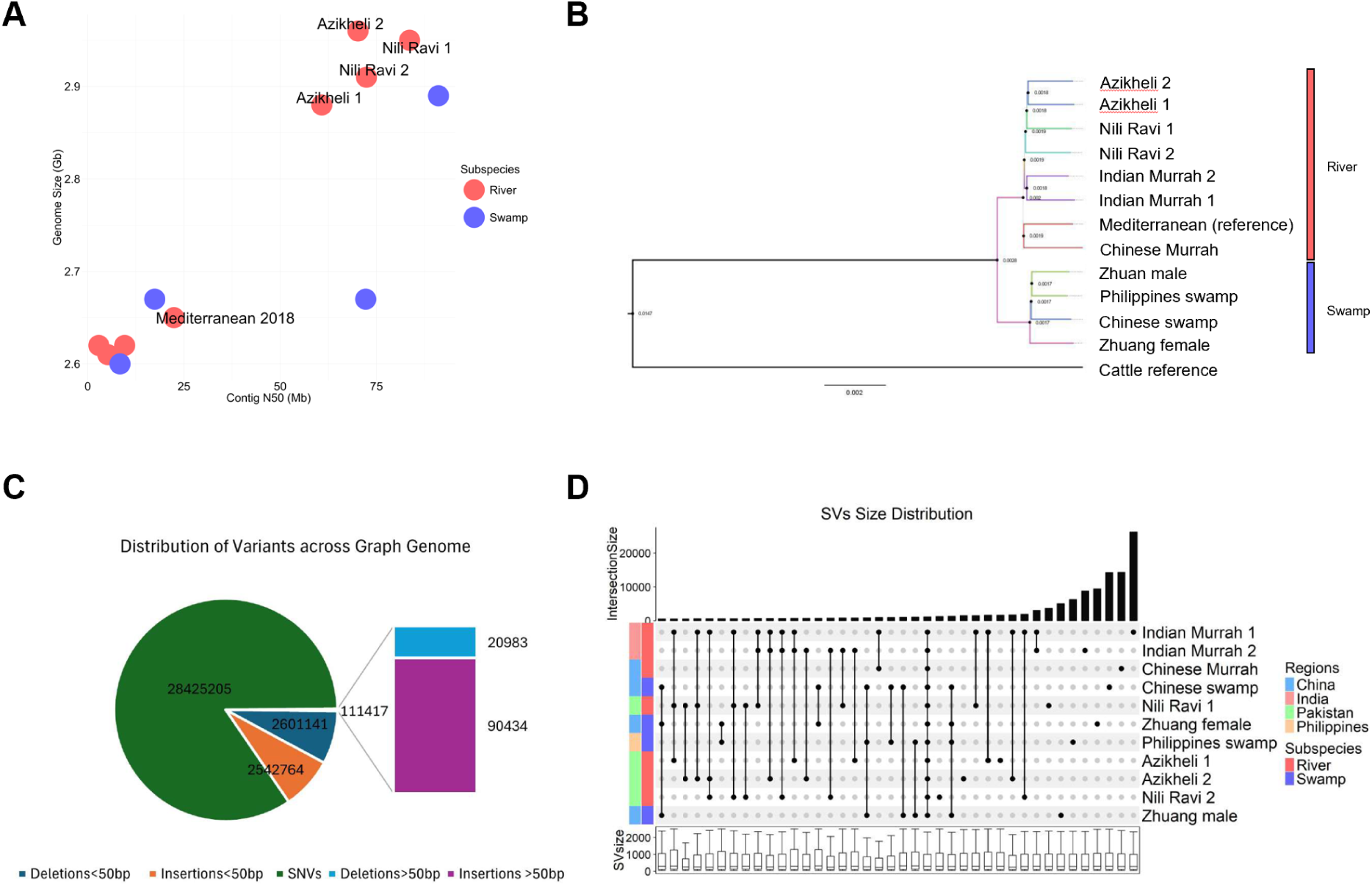
(A) Bubble plot of publicly available and newly generated genome assemblies, illustrating contig N50 on the x-axis and estimated genome size on the y-axis. Red bubbles represent river buffalo genomes, while blue bubbles denote swamp buffalo genome assemblies. The newly generated assemblies are highlighted as the top four red-labelled bubbles, surpassing the contiguity of the current Mediterranean reference genome (also labelled). (B) Phylogenetic tree showing the evolutionary relationships among the twelve water buffalo genome assemblies used in this study, including the newly generated haplotype resolved assemblies at the top and the cattle reference assembly (at the bottom) added as an outgroup. The tree was constructed based on the mash distances (distance matrix) method with 100 bootstrap replicates. Bootstrap values are displayed at the nodes. (C) Variant distribution across the water buffalo pangenome graph. The graph includes insertions and deletions (indels) <50 bp and structural variants (SVs) >=50 bp. (D) Upset plot of sets of SVs found across different assemblies. Each column represents a set of SVs with the points indicating in which assemblies the SVs were found. The bar graph along the top displays the number of SVs in the corresponding set. Only the 40 sets with the most SVs are shown.

**Table 1.** Genome assembly metrics for publicly available and newly generated haplotype resolved genome assemblies. All values were calculated using gfastats v1.3.6 [25] to ensure consistency. Originally reported metrics and assembly methods are compiled in Table 1 by [5], and found to be comparable.

| Assembly Name | Subspecies | Label | Genome<br>size(Gb) | Contigs | Contig<br>N50(Mb) | Contig<br>L50 | GC<br>content<br>% | Reference |
| --- | --- | --- | --- | --- | --- | --- | --- | --- |
| AZ0004_hap 1 | River | Azikheli 1 | 2.88 | 852 | 60.8 | 17 | 42.84 | Current study |
| AZ0004_hap 2 | River | Azikheli 2 | 2.96 | 582 | 70.2 | 16 | 43.22 | Current study |
| NR0003_hap 1 | River | Nili Ravi 1 | 2.95 | 510 | 83.6 | 14 | 43.18 | Current study |
| NR0003_hap 2 | River | Nili Ravi 2 | 2.91 | 674 | 72.4 | 16 | 43.01 | Current study |
| UOA_WB_1<br>(reference) | River | Mediterranean<br>2018 | 2.65 | 918 | 22.4 | 36 | 41.74 | [6] |
| BBCv1.0 | Swamp | Chinese<br>swamp | 2.67 | 8520 | 17.4 | 367 | 41.78 | [9] |
| NADDB_DH_1 | River | Indian Murrah<br>1 | 2.61 | 14366 | 5.22 | 1707 | 41.75 | [10] |
| NDDB_SH_1 | River | Indian Murrah<br>2 | 2.62 | 5265 | 9.59 | 636 | 41.85 | [10] |
| PCC_UOA_SB_1v2 | Swamp | Philippines<br>swamp | 2.89 | 297 | 91.1 | 29 | 42.66 | [5] |
| CUSA_SWP | Swamp | Zhuang<br>female | 2.60 | 9428 | 8.39 | 187 | 41.83 | [11] |
| CUSA_RVB | River | Chinese<br>Murrah | 2.62 | 11874 | 2.88 | 553 | 41.9 | [11] |
| Wang_2023 | Swamp | Zhuang male | 2.67 | 346 | 72.2 | 27 | 41.8 | [12] |

### Identification of structural variants based on pangenome graph

In order to create the water buffalo pangenome graph and identify structural variants (SVs), these four novel and seven public water buffalo assemblies (Table 1), were aligned to the reference genome (UOA_WB_1) using PanGenie’s minimap2 pipeline. The resultant pangenome graph contained a total of 32,821,198 variants, of which 2,542,764 and 2,601,141 were insertions and deletions less than 50 base pairs (indels) long, and 28,425,205 were single nucleotide variants (SNVs/SNPs). There were a further 111,352 structural variants (SVs), including 90,434 insertions and 20,983 deletions across the 24 autosomal chromosomes (Figure 1C). As shown in Figure 1D, the majority of structural variants (SVs) were found to be unique to individual assemblies. This observation is consistent with previous studies of cattle [43]. In total 1,347 SVs, spanning a total sum of 0.31Mb, were found specifically across all of the swamp genome assemblies, suggesting these SVs likely represent genomic segments specific to and fixed across this sub-species relative to river buffalo.

Emphasising the divergence of the two sub-species, among the top 40 sets of SVs shown in Figure 1D, only one set involved SVs shared across swamp and river assemblies – the set where the SVs were found in all of the non-reference assemblies - suggesting the variant is private to the reference assembly. Consequently, there is comparatively little SV sharing across sub- species.

In total, this buffalo pangenome graph contained an extra 147,865,364 bases in paths not present in the reference genome. The seven river assemblies exclusively contributed 73,672,251 bases, slightly higher than the exclusive 70,756,474 bases contributed by the fewer four swamp assemblies. Within the river assemblies, 38,960,389 bases were attributed to the novel Pakistani breeds (Azikheli 1, Azikheli 2, Nili Ravi 1, and Nili Ravi 2).

Consistent with good quality variant calling the transitions to transversions (Ti/Tv) ratio observed in our graph genome was 2.17, and comparable to the Ti/Tv ratio for whole-genome sequencing (WGS) data of *Bos taurus* and *Bubalus bubalus* in a previous study [16].

### Genetic variation observed across global water buffalo populations

We next sought to examine the frequency and segregation patterns of the variants identified in our pangenome across wider water buffalo populations. To do this we obtained whole genome sequencing (WGS) data from eight bioprojects (PRJNA633724, PRJEB39591, PRJNA547460, PRJCA001294, PRJNA350833, PRJNA1135737,PRJNA1057008, PRJNA633919) totalling 937 individuals. After filtering on depth of sequencing and sequencing approach, 711 were kept for downstream analyses, comprising 374 swamp and 337 river buffaloes. The geographic distribution of these samples are shown in Figure 2. The metadata details of the WGS cohort have been provided in Supplementary Table S1.

**Figure 2.**
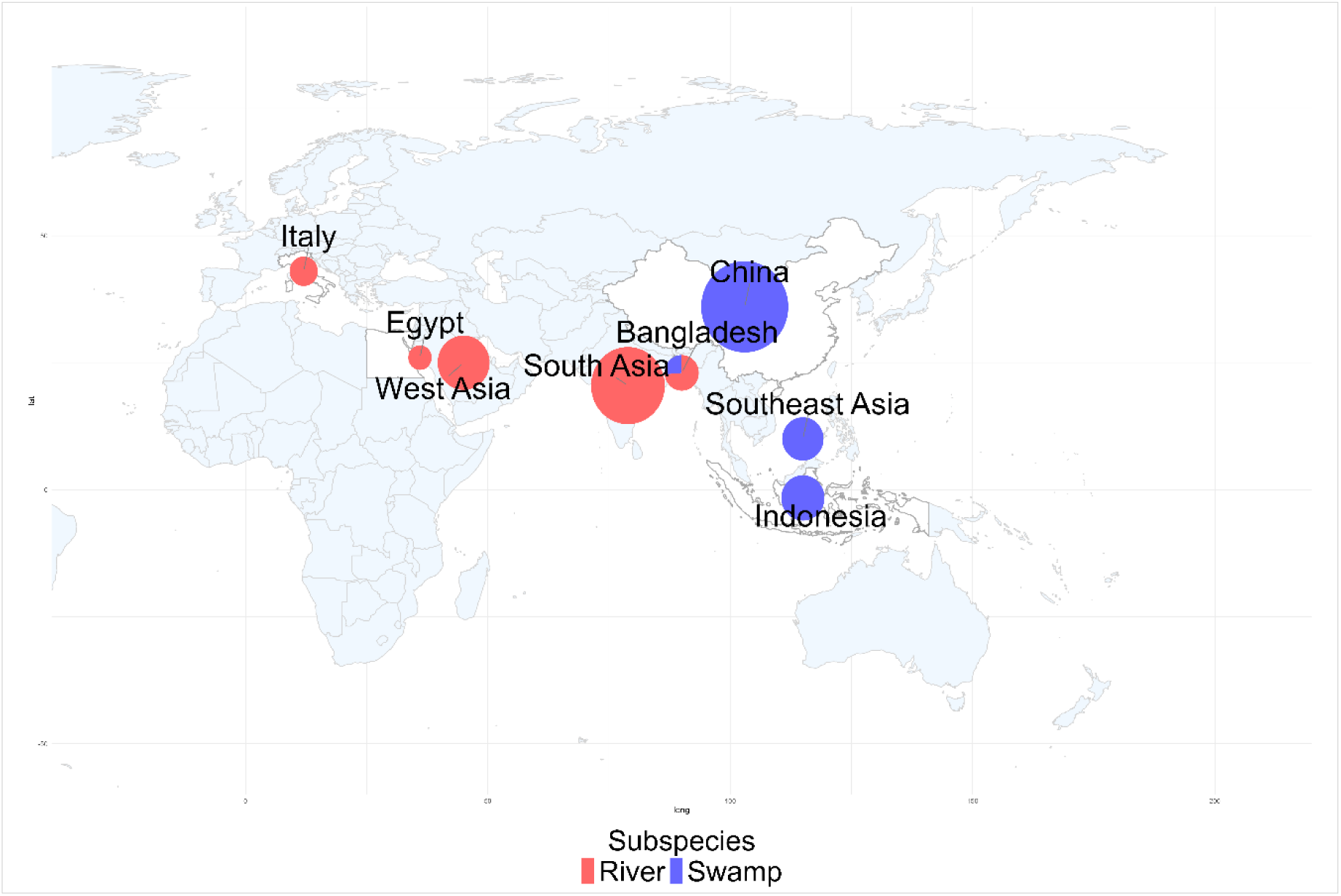
Geographic distribution of global water buffalo populations used in the study. The size of each pie corresponds to the relative sample size, while red and blue colours represent river and swamp buffalo subspecies, respectively.

Variants were called in each sample with respect to our newly generated reference graph genome (water buffalo pangenome) using PanGenie. Following stringent filtering for variants with a missing genotype rate >0.20 and minor allele frequency <0.05, 27,369,548 of 32,821,175 variants were retained for subsequent analysis.

For PCA and admixture analyses these samples were further filtered to remove closely related samples, leaving 403 individuals (175 river and 228 swamp buffalo) and following LD pruning, 678,504 biallelic SNVs were retained. As expected, genetic differentiation broadly reflects geography. The first principal component (PC1), accounting for 81.04% of the variance, and the admixture analysis at K=2, effectively separated the two buffalo subspecies, river and swamp (Supplementary Figs. S4, S5). Some evidence of admixture between the two sub-species is observed in the hybrid zone in Bangladesh, consistent with previous studies [17].

This cohort consequently provides a globally representative collection of water buffalo variant calls, both spanning the largest number of samples to date, and for the first time incorporating both short and longer variants. To enable reuse we have made this dataset available at (*Zenodo link on acceptance*).

### Concordance of graph and linear reference calls

Before undertaking downstream analyses with this dataset we wanted to address the open question of the relative advantages and disadvantages of graph based versus single reference based variant calling in water buffalo research. To begin to address this we examined the concordance of the PanGenie derived genotyping calls in a cohort of 81 samples to those derived from the traditional variant caller GATK. As shown in Figure 3A, for each sample, approximately 76% of SNVs were called by both callers, with the remaining variants relatively evenly split between those specific to GATK and those specific to PanGenie. As may be expected, these differences are disproportionately at repetitive regions of the genome. The majority (83%) of SNVs in non-repetitive regions being called by both genotypers. For non-SNVs, such as insertions and deletions, a higher proportion of variants were specific to PanGenie, including an extra 30% of variants being called in non-repetitive regions. In comparison GATK only called an extra 20% of non-SNV calls in these regions. These results are broadly consistent with the idea that graph callers are potentially better able to detect non-SNV calls than traditional genotyping tools. However, an important disadvantage of callers such as PanGenie is their limitation to only calling genotypes at variants represented in the genome graph. This is emphasised when examining the allele frequency of variants specific to one or other caller (Figure 3B). GATK specific calls are comparatively enriched with those with a low allele frequency. This is consistent with PanGenie missing these rarer variants due to their lower frequency and therefore their lower probability of being represented in the graph. No substantial difference in the proportion of variants out of Hardy-Weinberg equilibrium was observed between the sets of variants specific to each caller (Supplementary Fig. S6). However, a difference in transition/transversion (Ti/Tv) ratios was observed, with the PanGenie specific variant calls generally having a lower ratio (Supplementary Fig. S7), associated with putatively more false positives. This likely in part reflects that the GATK calls were filtered based on metric cutoffs guided by Ti/Tv ratios [16]. On average the Ti/Tv ratio of the 711 PanGenie calls in each individual was 2.18 (Supplementary Table S3). Notably this is higher than what was observed in the original PanGenie paper where a Ti/Tv ratio of around 2.01 was observed for human assemblies [26]. Consequently the optimum variant caller will likely depend on the planned downstream analyses. Analyses such as the study of selective sweeps or genome-wide association studies where low frequency variants are often filtered out will benefit less from the advantages of GATK, particularly given its longer run time. However, studies where it is necessary to detect private or low frequency variants and reduce false positive SNV rates, for example the study of mutation rates, will be at a disadvantage if graph-based callers such as PanGenie are used.

**Figure 3.**
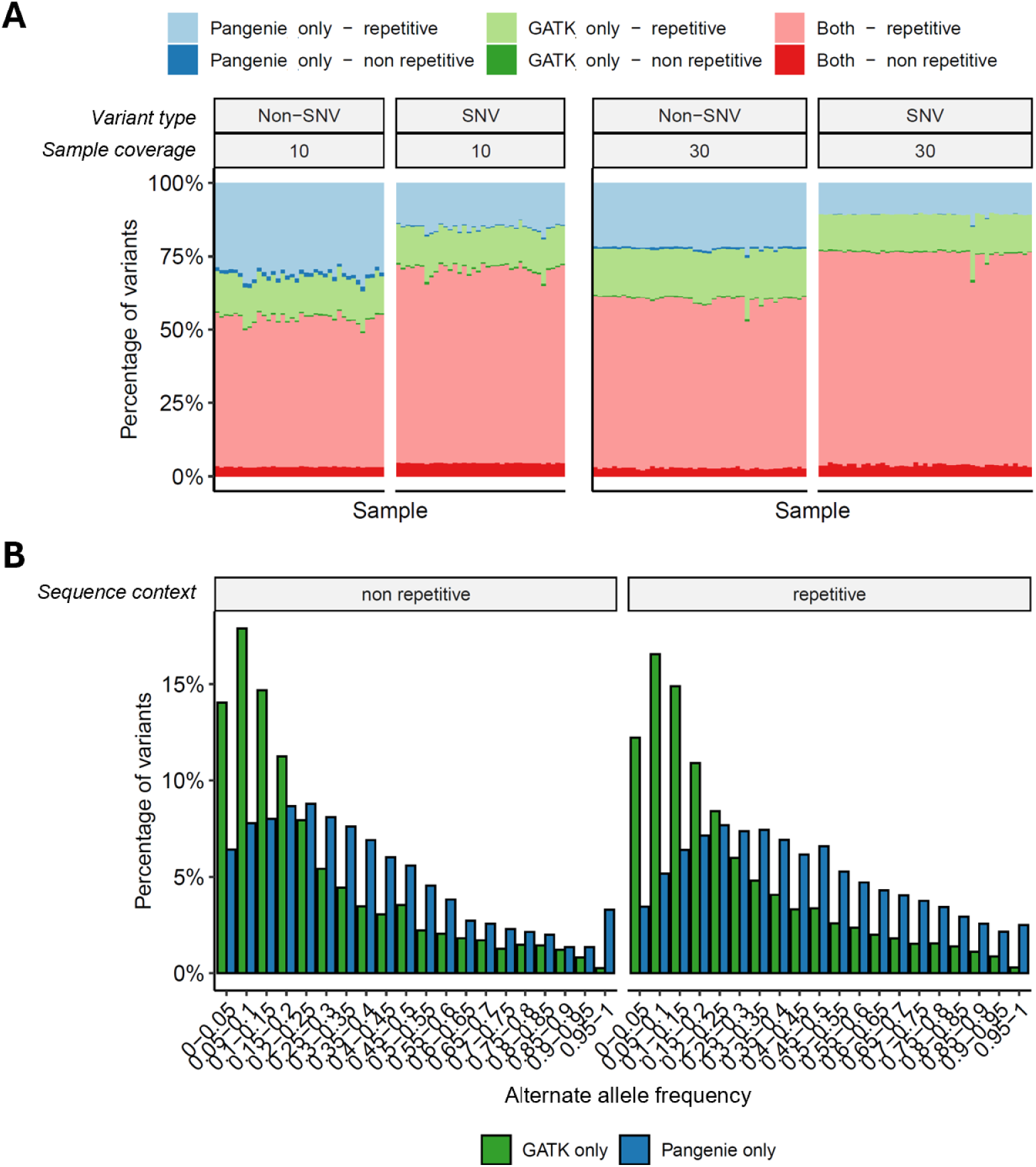
The overlap of variants called by GATK and PanGenie across 81 river buffalo samples. In each plot each column corresponds to a sample and the Y axis indicates the proportion of variants called by both variant callers (red) or only by PanGenie (blue) or GATK (green). Intensity of colour indicates the variants’ sequence context (found in repetitive or non-repetitive sequence contexts). Panels are further broken down by variant type (SNV or non-SNV) and the approximate coverage of the samples (10x or 30x sequencing coverage). (B) The allele frequencies among the samples of variants specifically called only by either GATK or PanGenie in the high coverage (30x) samples. The results are broken down according to whether variants are found in repetitive regions.

### Selective sweeps in water buffalo

We next explored the utility of graph genomics approaches to inform the identification of functional genes and variants under selection in water buffalo. To do this we restricted our larger cohort to populations with at least six unrelated individuals. This resulted in a set of 282 samples spanning 26 distinct populations (Supplementary Table S2), consisting of 15 swamp buffalo and 10 river buffalo groups. The integrated haplotype score (iHS) and number of segregating sites by length (nSL) statistics were then calculated within each group to identify sites of potential positive selection. In total 1960 genes were detected under a putative selective sweep peak for one or other metric, with 249 genes identified by both (1065 only detected by iHS and 646 by nSL, Supplementary Table S4). To explore the significance of these genes, we conducted gene set enrichment analyses using FUMA [44], focusing on enrichment among genes linked to traits in human genome-wide association studies (GWAS). Intriguingly, a range of relevant phenotypes were preferentially associated with the genes under putative selective sweep peaks (Figure 4). For iHS these ranged from obesity-related traits and adult body size, to coat colour and immune-relevant phenotypes such as mosquito bite size. Furthermore, nSL highlighted additional behavioral phenotypes from anxiety and stress-related disorders, to dental health indicators such as smooth-surface caries (Supplementary Table S5). These results consequently provide insights into the target phenotypes and underlying genes under selection in water buffalo. All population-level iHS and nSL scores can be viewed along the genome alongside other annotations including XP-EHH and XP-CLR scores from a previous study [16] at our Bovine Omics Atlas browser (www.bomabrowser.com/jbrowse/index.html?data=BOMA/v1.0), enabling exploration of selection signals across water buffalo populations.

**Figure 4.**
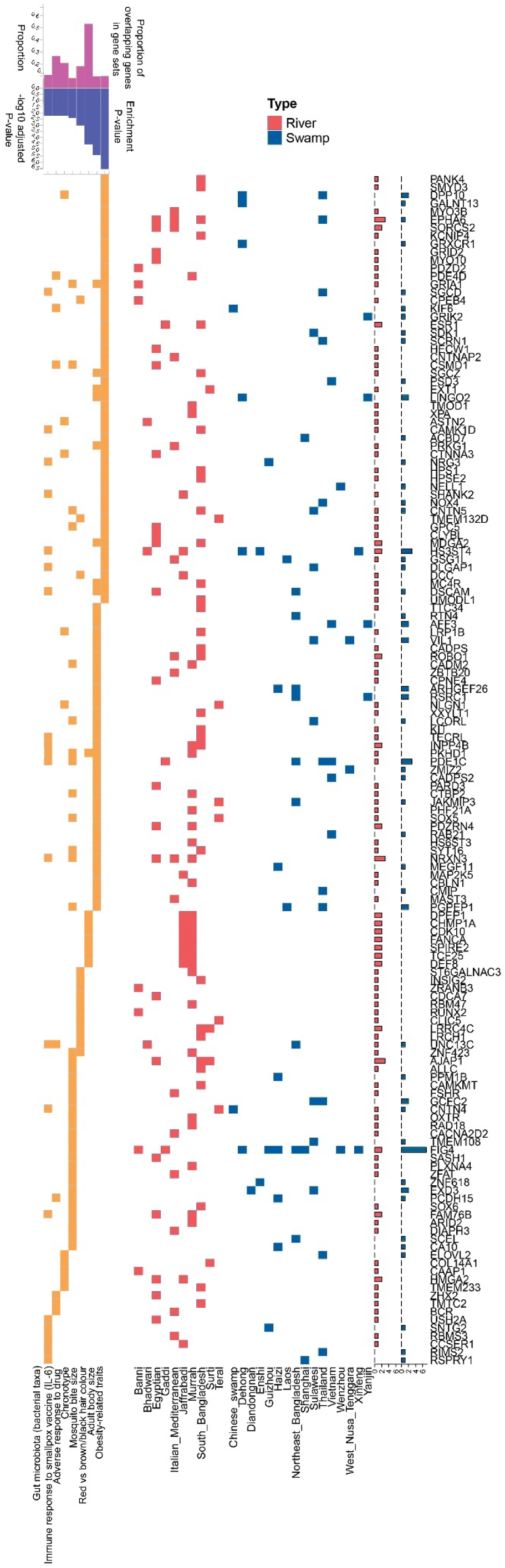
Enrichment analysis of the genes under peaks identified in the iHS analysis. The x-axis of the first panel shows human GWAS traits enriched among the genes falling under iHS peaks (as identified by FUMA), the x axis of the second panel breeds in which the corresponding selective sweeps are observed. The y-axis lists the genes within the respective gene sets and peaks, with the boxes indicating with which trait and selective sweep in which breed they are associated with. Enrichment P-values and adjusted P-values (expressed as -log₁₀ values) are shown at the top left to indicate the enrichment of terms (only the top eight most significant terms are shown). The bar graphs on the left show the total number of breeds in which a putative selective sweep peak that intersected the corresponding gene was observed.

More specifically, 53 genes were associated with the “obesity-related traits” term (P-value of 6.09 × 10⁻¹¹ and an adjusted P-value of 2.69 × 10⁻^7^) in the iHS analysis including *MC4R* in South Bangladeshi river buffalo, mutations of which are the commonest form of monogenic obesity in humans [45]. Likewise *LCORL*, under putative selection in Sulawesi swamp buffalo and among the 46 genes linked to body size, has been linked to birth weight and growth in various cattle studies [46, 47], suggesting this gene has been targeted by domestication across bovids. Intriguingly 14 genes were linked to the “mosquito bite size” term and six to the “immune response to smallpox vaccine” term (Figure 4) suggesting both artificial selection for production traits as well as natural selection for immune traits are key drivers of selective sweeps across the water buffalo populations.

### Identifying candidate larger variants linked to selective sweeps

A key advantage of graph genomics approaches is the ability to assay larger variants that may drive variation in phenotypes and traits, but that may have been missed in traditional approaches focused on SNPs. To investigate putative adaptive variants affecting coding regions, we annotated variants using SnpEff v5.2f (2025-02-07) [42] and SnpSift v5.2f [48]. Prior to annotation, multiallelic variants were normalized by splitting them into separate biallelic entries, resulting in 6,159,686 indels, 28,669,966 SNVs, and 160,921 SVs entries. Within putative selective sweep regions we identified 208,862 indels, 997,500 SNVs and 6,748 SVs. Notably an enrichment of HIGH impact SVs, indels and SNVs were observed within selective sweep regions (Figure 5A, Supplementary Table S6), with 50-80% more variants in these areas having a HIGH impact compared to genome-wide. Among the high impact variants in selective sweep regions only 20% were SNVs, with the remainder being SVs and indels, suggesting high impact larger variants may underlie putative selective sweeps.

**Figure 5.**
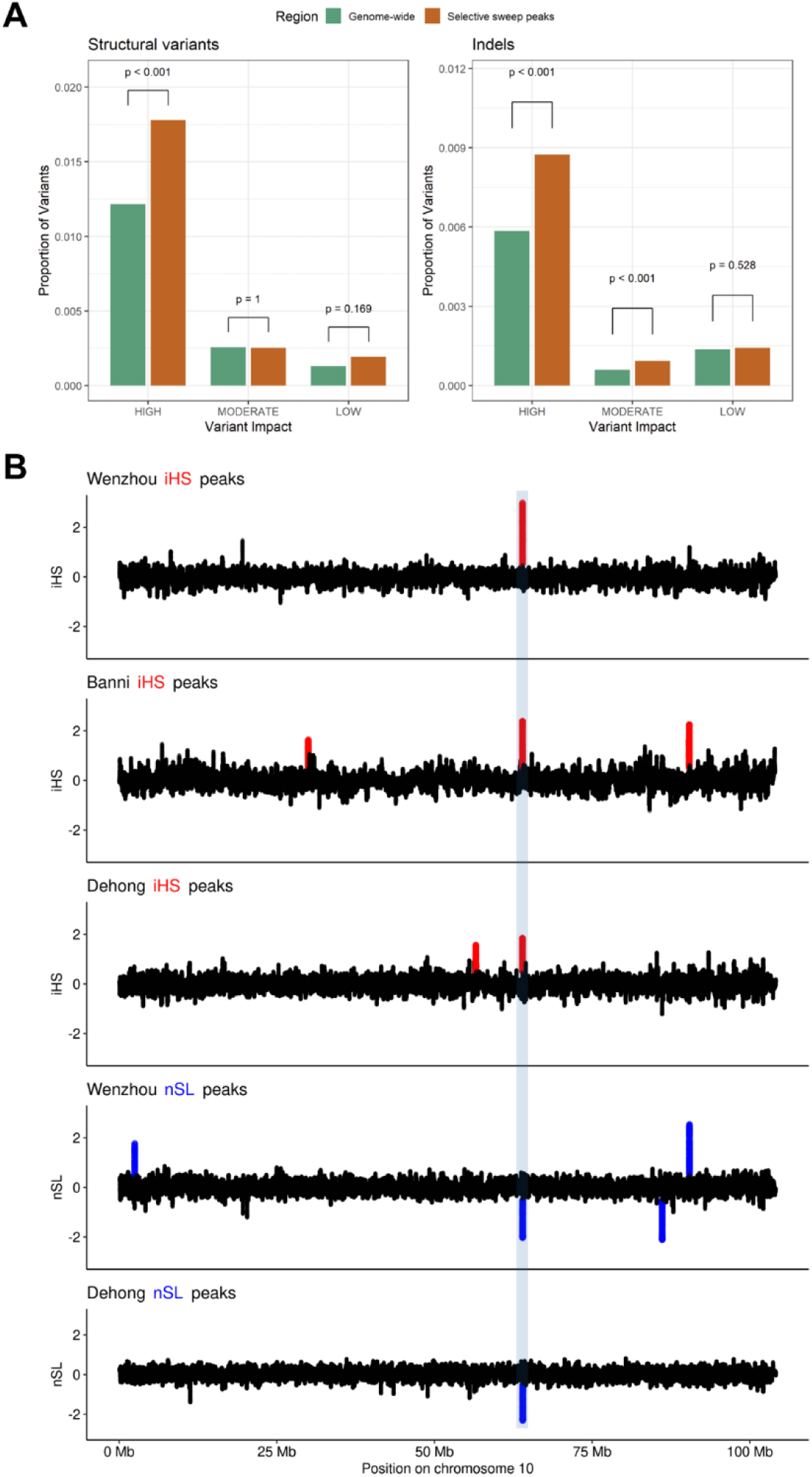
(A) Enrichment of high impact SVs and indels in selective sweep peaks. The y-axis shows the proportion of variants in each category (genome-wide or selective sweep peaks) that are in each impact class (HIGH, MODERATE and LOW. Due to the disproportionate size of their bars the MODIFIER class is not shown). Two-sided Fishers exact P values are shown above the bars of the difference between categories of the proportions of variants in the corresponding impact class. (B) Example colocalization of selective sweep peaks observed across populations and metrics at the *FIG4* locus. Called peaks are indicated by red (iHS) or blue (nSL) points with the respective buffalo population indicated above the plot.

In total 965 genes were affected by HIGH impact variants falling in a putative selective sweep peak. Among the indels and large structural variants with a predicted HIGH or MODERATE impact several also exhibited pronounced differences in alternate allele frequencies (aAF) between populations consistent with putative selection, including those linked to production, fertility, immunity or adaptability traits. The detailed summary of genes associated with HIGH consequences has been provided in (Supplementary Table S7).

Evidence for a strong selective sweep signal was observed exclusively in river buffalo populations at the *PGRMC2* (Progesterone receptor membrane component 2) locus on chromosome 17 (Supplementary Fig. S8). The *PGRMC2* gene is associated with fertility and production-related traits in bovines [49] and is additionally expressed in bovine mammary tissues during lactation in dairy cattle [50, 51].

A potential HIGH-impact 11 base pair (bp) insertion was observed within the coding region of this gene at position 17:43,563,676. This insertion occurs at cDNA position c.294_295 in transcript XM_006067309.2, resulting in a frameshift starting at amino acid position 99. This frameshift introduces a premature stop codon downstream, disrupting the C-terminal cytoplasmic domain of *PGRMC2* [52]. Consistent with the observed difference in selection between the sub-species the average alternate allele frequency (aAF) of this variant was 91% in swamp buffalo, compared to the river buffalo populations which exhibited an average alternate allele frequency of 32%. Given the uterus’s pivotal role as a target for progesterone (P4) responses, disruptions in *PGRMC2* function are potentially likely to impair uterine function and fertility [53]. This is corroborated by epidemiological studies in humans [52, 54, 55] and livestock [52, 56], as well as genetic research in rodents [52, 55], which link low conception rates to inadequate progesterone levels and subsequent uterine dysfunction. In cattle, a similar region on chromosome BTA17 (spanning 29 Mb to 34 Mb) is also strongly associated with milk fatty acid composition [51].

Selective sweeps spanning two neighbouring genes, non-SMC conodensin I complex subunit G (*NCAPG*) and ligand dependent nuclear receptor corepressor-like (*LCORL*) were identified on chromosome 7 (Supplementary Fig. S9). *NCAPG* encodes a subunit of the condensing 1 protein involved in chromatin condensation during replication and found to be associated in modulating fetal growth in cattle [57, 58]. *LCORL* is thought to be a transcription factor that may function during spermatogenesis in the testes and may have associated roles in height, growth and withers in equines [58, 59]. Numerous studies have shown genetic variation at the *LCORL- NCAPG* locus is strongly associated with body size and growth traits in beef cattle [47]. In our study these genes were found to be under strong selective sweep in Sulawesi swamp buffalo (Supplementary Fig. S9). The *LCORL* gene contains a MODERATE impact deletion resulting in the loss of three alanine residues (p.Ala21_Ala23del), within exon 1 of the gene. This particular variant has the highest alternate allele frequency in the Sulawesi swamp population (AF of 0.91) consistent with selection in this group. Importantly variation in this gene has been linked to production. Perhaps most notably, an *LCORL* frameshift variant has been linked to a range of production and morphology traits in cattle [47].

A large 14kb HIGH impact deletion, detected at position 10:63987503-64002148 and predicted to lead to a transcript ablation of the *FIG4* gene, fell under a selective sweep peak detected across a large number of swamp buffalo genomes (Figure 5). *FIG4* is crucial for neuronal and muscular functions via phosphoinositide signalling pathways [60].

A mucin-3A-like gene (LOC102408548) falls under a selective sweep peak detected in the Shanghai, Northeast Bangladesh, Thailand and Haizi buffalo populations (Supplementary Fig. S10). Within this gene, we detected a MODERATE impact disruptive in frame insertion of 390bp specific to swamp buffalo, for which the reference allele is fixed across river buffalo populations. In contrast this insertion shows a strikingly elevated alternate allele frequency in the corresponding Shanghai (frequency of 1), Northeast Bangladesh (0.5), Haizi (0.68) and Thailand (0.70) buffalo populations. Mucin-3A *(MUC3A*), is an epithelial glycoprotein found along the mucosal lining and plays protective role against infectious agents and particles by providing lubrication and maintaining integrity https://www.uniprot.org/uniprotkb/Q02505/entry#function [61]. The elevated allele frequency in relevant buffalo populations underscores the potential relevance of this SV.

These findings highlight these genes as example prime candidates for further investigation into their role in buffalo adaptation and resilience, with potential implications for conservation and selective breeding programs.

## Discussion

In this study, we present the first high-quality genome assemblies for Pakistani water buffalo breeds, integrate these assemblies into a comprehensive water buffalo pangenome, perform the largest assessment of global water buffalo variation (including structural variants) to date, and use these data to explore how positive selection targeting larger variation may be driving important buffalo phenotypes. We have made these data publicly available, including via a genome browser (www.bomabrowser.com/jbrowse/index.html?data=BOMA/v1.0) enabling users to browse the selective sweep data across the genome and relative to other annotations.

Our newly generated Pakistani assemblies rank among the most contiguous water buffalo genomes available, and are the most contiguous for river buffalo in terms of contig N50, exceeding even the current reference genome. In this study we chose to focus on creating dual assemblies, i.e. two pseudo-haplotype genomes per animal, due to our primary focus on assaying structural variants. Although many previous studies, including the generation of the current reference, have produced collapsed assemblies, i.e. one genome for an individual, this has the disadvantage of reducing the number of variants that can be detected. For example, at best, only one allele can be integrated at heterozygote sites. However, using our publicly released data it would be possible to produce collapsed assemblies with even higher metrics, with contig N50s of 90.6Mb (Azikheli) and 83.6Mb (Nili Ravi) expected, and consequently comparable to the Philippine swamp buffalo assembly, the currently most contiguous collapsed assembly.

One metric on which our assemblies rank lower is scaffold N50, largely because we chose not to invest resources in scaffolding. Given that graph pangenomes focus on aligning orthologous contigs across assemblies and identifying shared or unique sequences, extensive scaffolding offers comparatively little added value for pangenome construction.

Our integrated pangenome revealed over 140 Mb of non-reference paths. Although fewer swamp buffalo assemblies were included, these contributed approximately the same amount of novel sequence as the river buffalo assemblies. This likely partly reflects the fact that the main reference genome is river-derived, so more of the river-specific variation is already captured. Additionally, most structural variants appear to be sub-species–specific, consistent with limited introgression—only at the Bangladesh interface is there evidence of any appreciable gene flow between river and swamp buffalo lineages.

One potential hurdle to implementing water buffalo pangenomics has been the divergent karyotypes (2n = 50 vs. 2n = 48). Our approach, in which we split swamp buffalo chromosome 1 at its fusion point to align with the river buffalo karyotype, illustrates a straightforward solution to incorporate both sub-species into the same pangenome. Future studies may benefit from tools that automate such chromosomal splits, allowing for broader applications of graph genomics across diverse buffalo populations.

A comparison of GATK [62] (reference-based) and PanGenie [26] (graph-based) genotyping highlights the complementary strengths and limitations of each. Graph-based caller approaches are especially well suited to identifying structural variants—one of the central aims of this work— while reference-based methods, such as GATK or DeepVariant [63], frequently excel in *de novo* detection of novel SNPs and often produce fewer lower-frequency false positives. In practice, the optimal choice of method thus depends on specific research goals: if discovery of large, functionally important variants is paramount, graph-based approaches may prove particularly advantageous; for high-confidence, short variant calls, traditional workflows remain valuable. In certain cases, combing both, i.e. shorter variant calls from single reference methods and larger calls from graph-based methods may be optimal.

Applying selective sweep analyses to one of the largest water buffalo genomic datasets assembled so far enabled us to pinpoint hundreds of genes putatively under positive selection. Over 200 genes were identified by both of the statistical methods adopted, underscoring the robustness of these signals. Notably, these genes were frequently enriched for traits related to growth, size, and immune response—phenotypes likely under strong natural and artificial selection in water buffalo. Crucially, we identified multiple candidate functional larger variants in these regions, which may go undetected by single reference-based, SNP-centric approaches. These findings not only highlight the potential of graph-based genomics for discovering new selection signals but also open avenues to integrate such structural variants into breeding programs aimed at enhancing productivity, disease resistance, and other economically important traits in water buffalo.

Although our focus in this study was characterising the potential relevance of larger variants to selective sweeps, the variant call set from this study would have utility to a diverse range of other projects. From acting as a reference panel enabling the imputation of both short and long variants, to studying the occurrence of compound heterozygote loss-of-function variants.

Overall, our work provides important novel resources and insights into how graph genomics can accelerate our understanding of structural variation and its role in driving phenotypic diversity across a species characterized by multiple karyotypes and a complex domestication history.

## Supplementary Figures

**Supplementary Fig. S1.** The Nili Ravi heifer from the Punjab province of Pakistan selected for the whole genome assembly.

**Supplementary Fig. S2.** The female Azikheli river buffalo from Swat district of Pakistan sampled for the whole genome assembly.

**Supplementary Fig. S3.** The cross-validation error plot depicting CV values across different values of K, with K=6 highlighted with the lowest CV error.

**Supplementary Fig. S4.** Principal Component Analysis (PCA) plot using the graph genome as a reference reveals clear geographical clustering both between and within the river and swamp buffalo populations.

**Supplementary Fig. S5.** Admixture plot for different K values ranging from K=2 to K=6 effectively representing the significant zones of hybridization between the populations.

**Supplementary Fig. S6.** The distribution of Hardy-Weinberg equilibrium (HWE) P values of variants specifically called by GATK (green) or PanGenie (blue). Variants are broken down into those falling or not falling into repetitive regions and the genome-wide significance threshold of 5x10^-8^ is indicated by grey dashed lines. Neither variant caller shows a large number of variants above this threshold.

**Supplementary Fig. S7.** The distributions of per sample transition/transversion ratios of the SNVs called by both variant callers (red) or by only GATK (green) or PanGenie (blue). Results are broken down by the approximate coverage of the samples and whether the variant falls within a repetitive region.

**Supplementary Fig. S8.** The genome-wide selective sweep peaks within the *PGRMC2* gene at the chromosome 17 locus identified by iHS analysis, revealed a pronounced selective sweep signal in river buffalo populations but not in swamp buffalo populations.

**Supplementary Fig. S9.** The putative Sulawesi selective sweep spanning two candidate genes

*NCAPG* and *LCORL* on chromosome 7.

**Supplementary Fig. S10.** The prominent selective peaks within the *MUC3A* gene, specific to the swamp buffalo population.

## Supplementary Tables

**Supplementary Table S1.** The data accession numbers and relevant information for each whole- genome sequencing (WGS) biosamples used to capture genetic variation across global water buffalo populations.

**Supplementary Table S2.** The details of biosamples accessions and grouping information of population groups selected for the selective sweeps analysis, in which the population with least six unrelated individuals per group was included.

**Supplementary Table S3.** Summary statistics of PanGenie genotyped WGS cohort.

**Supplementary Table S4.** The list of genes identified in potential selective sweeps, categorizing those exclusive to the iHS and nSL metrics as well as those identified by both of these.

**Supplementary Table S5.** The list of FUMA enrichment analysis of genes involved in selective sweeps identified by the iHS and nSL metric, detailing gene sets, associated trait terms and p- values.

**Supplementary Table S6.** The counts of HIGH,MODERATE,LOW and MODIFIER impact variants, categorized into SNVs, Indels, and large SVs, across genome-wide autosomal regions and selective sweep regions.

**Supplementary Table S7.** The list of genes along with descriptions of their roles that are impacted by HIGH-impact variant consequences within regions identified as undergoing selection.

## Data availability

The PacBio HiFi sequencing data generated in this study, including raw FASTQ reads and haplotype-resolved genome assemblies for *Nili Ravi* and *Azikheli* water buffalo, have been deposited in the European Nucleotide Archive (ENA) under BioProject accession PRJEB86148 (study accession: ERP169523). The corresponding BioSample accessions are SAMEA117759395 (*Nili Ravi*) and SAMEA117759394 (*Azikheli*). Each individual has two haplotype-resolved assemblies, available under the following analysis accessions: ERZ26095406 and ERZ26095324 (*Nili Ravi*); ERZ26095169 and ERZ26095065 (*Azikheli*).

The genome Assemblies UOA_WB_1 (GCF_003121395.1), NDDB_DH_1(GCA_019923925.1), NDDB_SH_1 (GCF_019923935.1) and PCC_UOA_SB_1v2 (GCF_029407905.1) were downloaded from NCBI and CUSA_SWP (GWHAAJZ00000000) and CUSA_RVB (GWHAAKA00000000) were accessed and downloaded from NGDC. The genome assembly Wang_2023 was downloaded from figshare as documented in[12]. The genome assembly BBCv1.0 was obtained from the Sequence Archive CNSA under the project accession CNP0000797, which can be accessed at https://ftp.cngb.org/pub/CNSA/data5/CNP0000797/CNS0152939/CNA0007311/.

## Acknowledgements

We gratefully acknowledge the Commonwealth Scholarship Commission for funding this research through the Commonwealth Split-site PhD Scholarship Program, which supported FA’s one-year visit to the Roslin Institute and enabled the successful execution of the core components of this study. We also thank Dr. Rahimullah, Veterinary Officer at the “Azikheli Buffalo Improvement and Conservation Farm, Charbagh, Swat”, Pakistan, for his assistance with sample collection. This work was further supported by grants BB/T019468/1 and BBS/E/RL/230001A from the UK’s BBSRC funding council.

## Author Contributions

The initial idea for this work was conceived by JP, FA and SJ. The animal blood sampling and DNA extraction was performed by MA, MM and SM while RO provided technical support for the initial quality control of samples. The formal analysis of assembly preparation, graph genome construction, and downstream analysis, was conducted by FA with coding support from SJ and AT, under the supervision and validation of SJ and JP. The selective sweep and concordance analysis was carried out by JP and FA. The initial draft of the manuscript was jointly written by FA and JP, with revisions provided by SJ and AT.

## Competing Interests

The authors have no conflicts of interest to disclose.

| Contributions | Authors |
| --- | --- |
| Conceptualization | JP,FA,SJ |
| Data Curation | MM, MA,SM,RO |
| Formal Analysis | FA |
| Software | SJ, JP, AT |
| Supervision | JP, SJ |
| Validation | SJ, JP |
| Writing Original Draft | FA, JP |
| Writing Review and Editing | SJ, AT, JP |

