## Supplementary Figures 1-10 for "A comprehensive water buffalo pangenome reveals extensive structural variation linked to population specific signatures of selection"

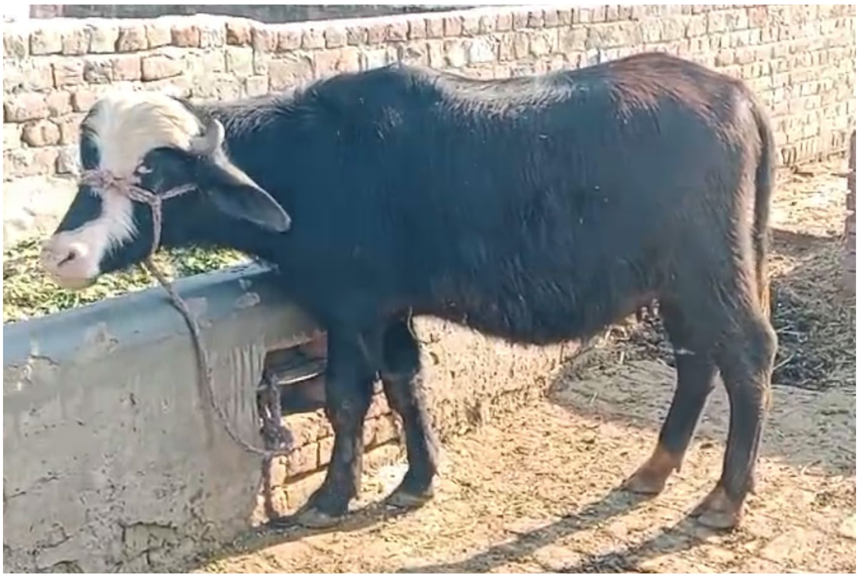

**Supplementary Fig. S1**

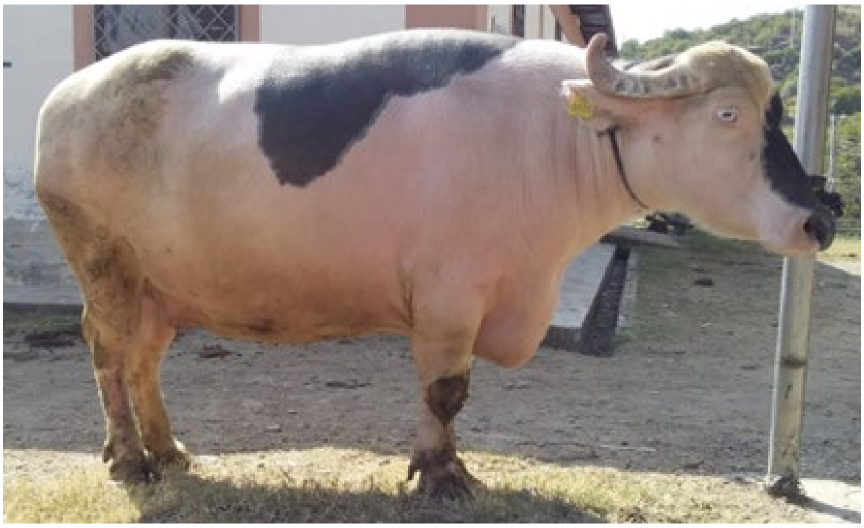

**Supplementary Fig. S2**

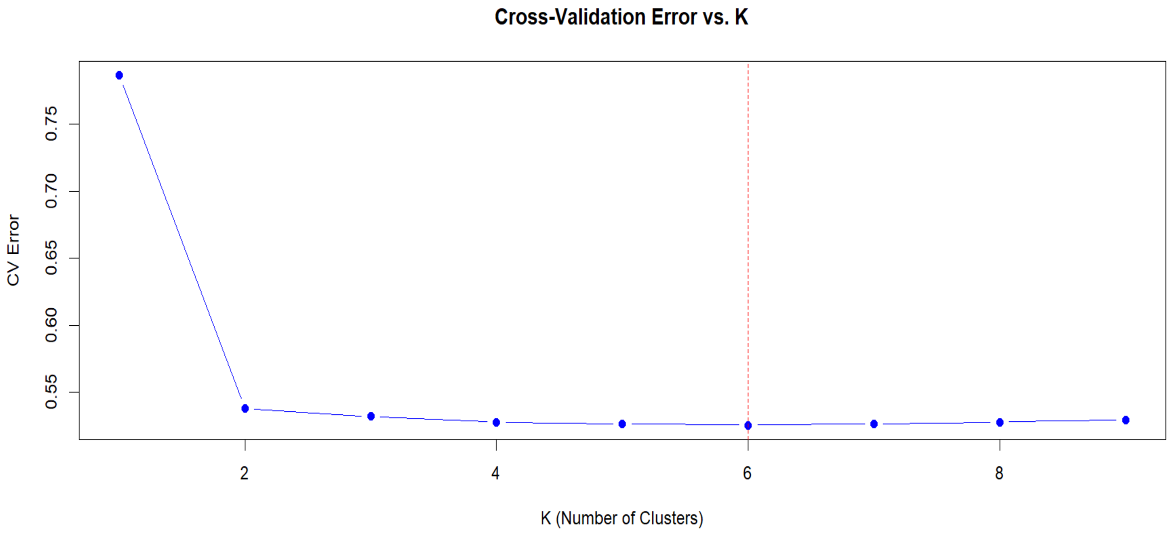


**Supplementary Fig. S3**

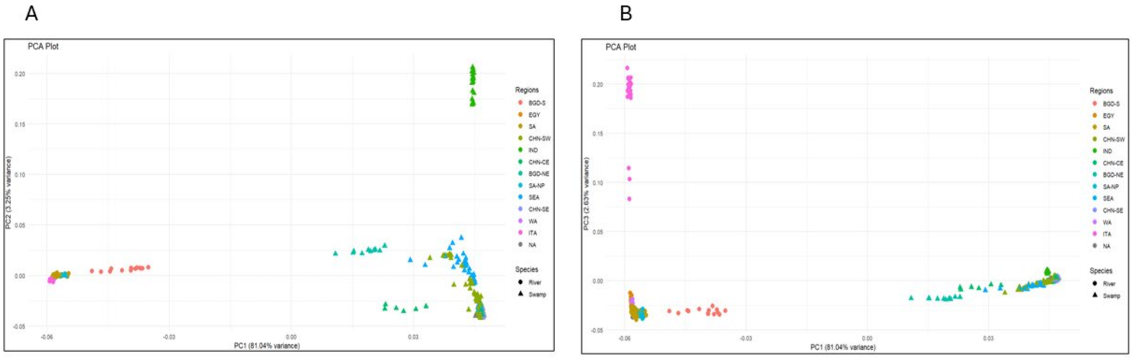


**Supplementary Fig. S4**

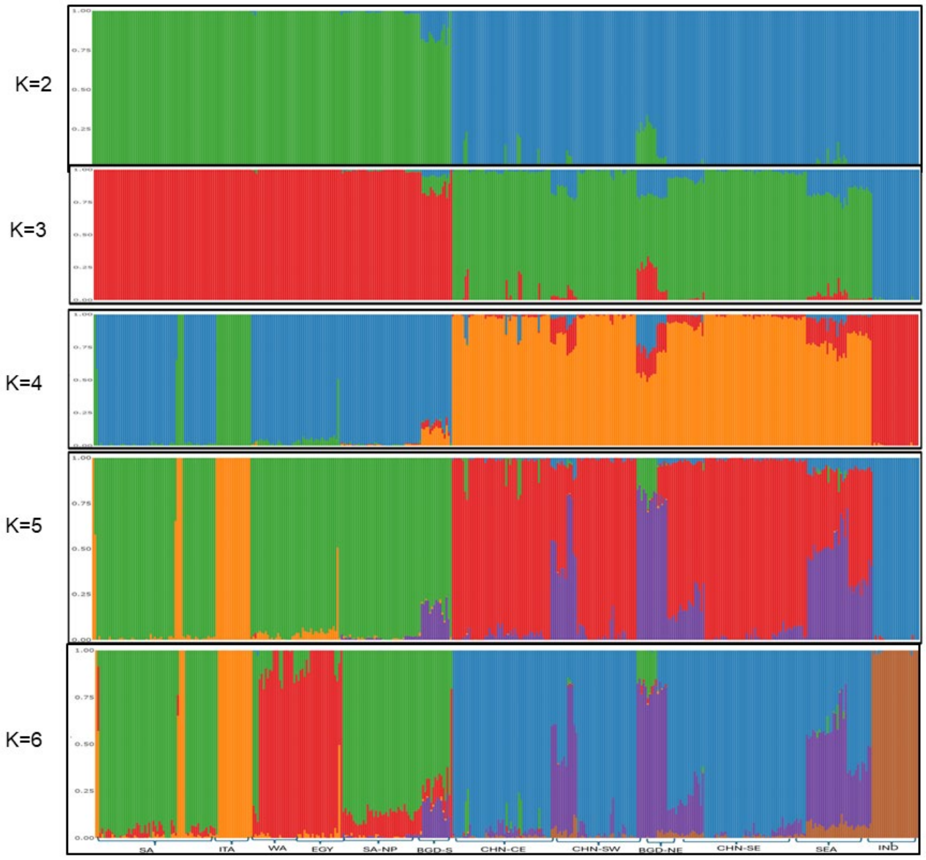


**Supplementary Fig. S5**

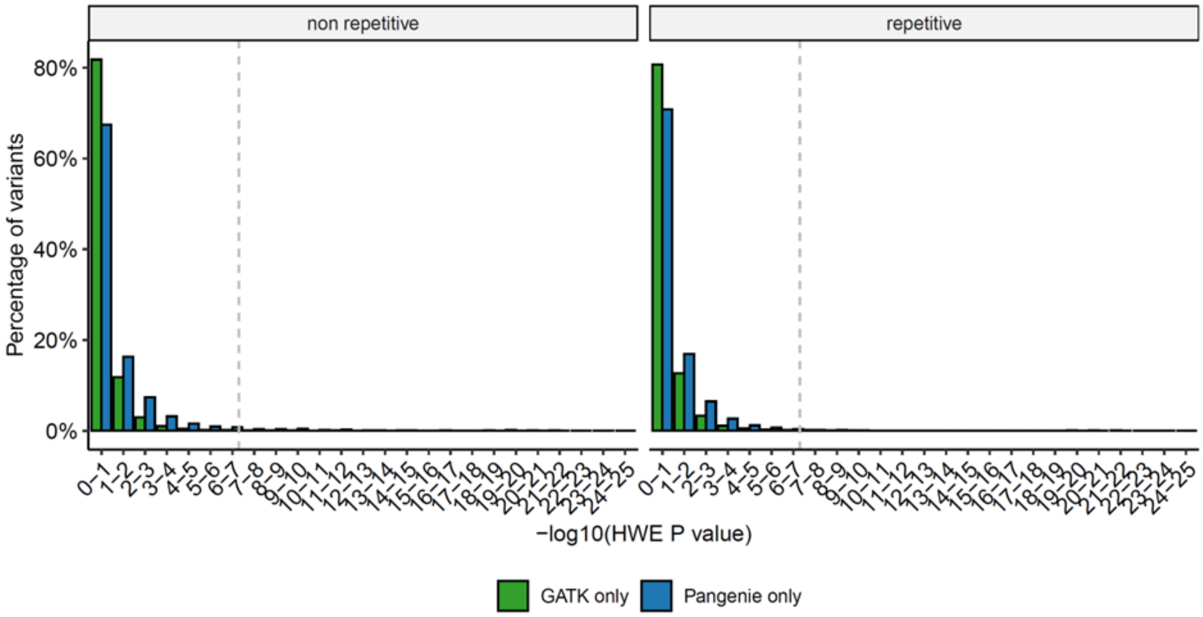


**Supplementary Fig. S6**

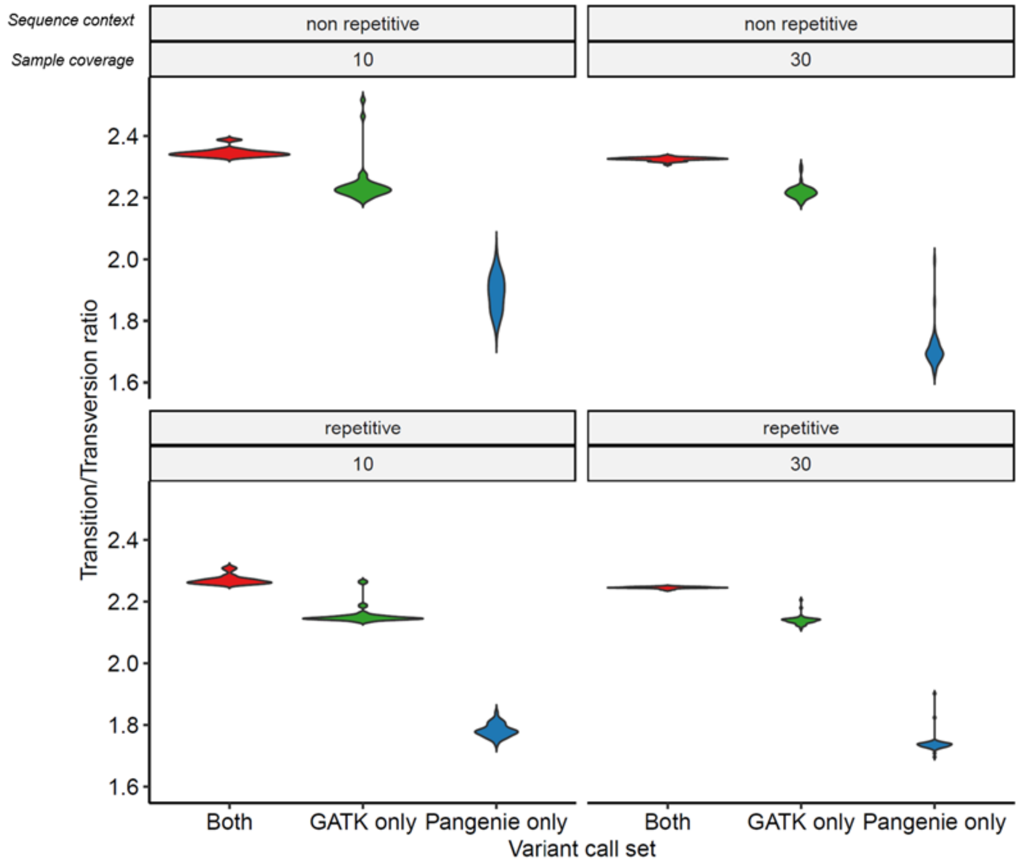


**Supplementary Fig. S7**

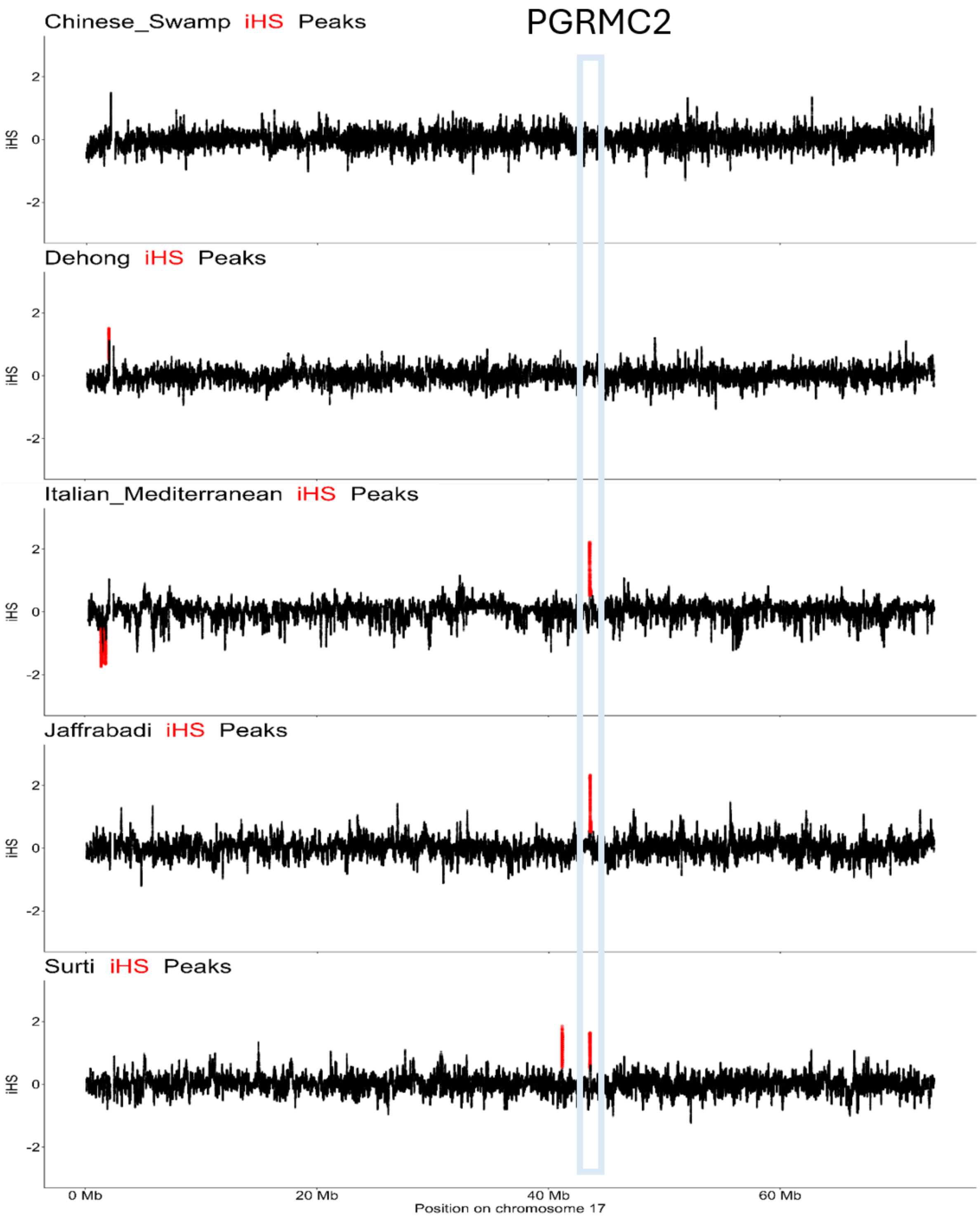


**Supplementary Fig. S8**

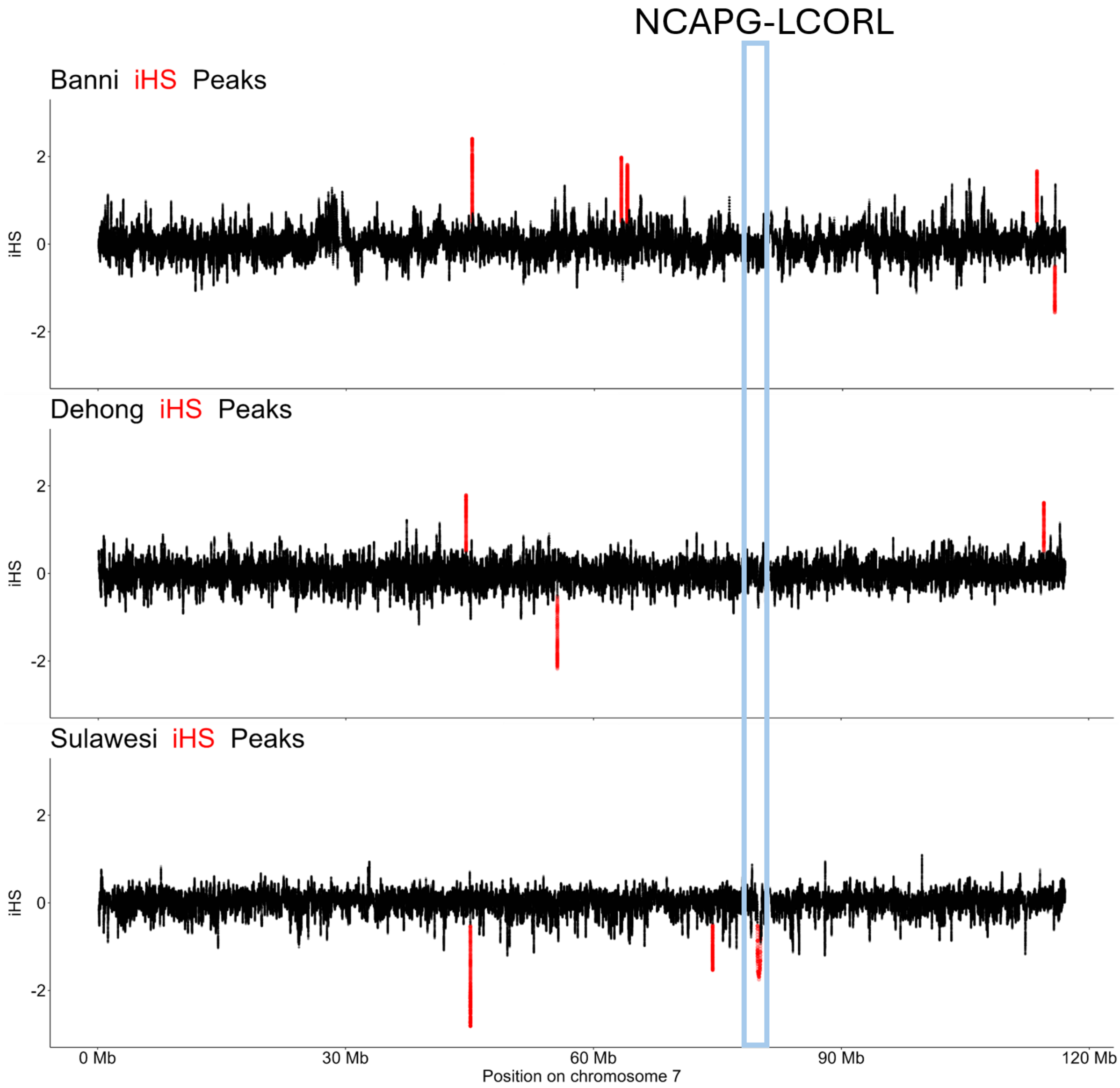


**Supplementary Fig. S9**

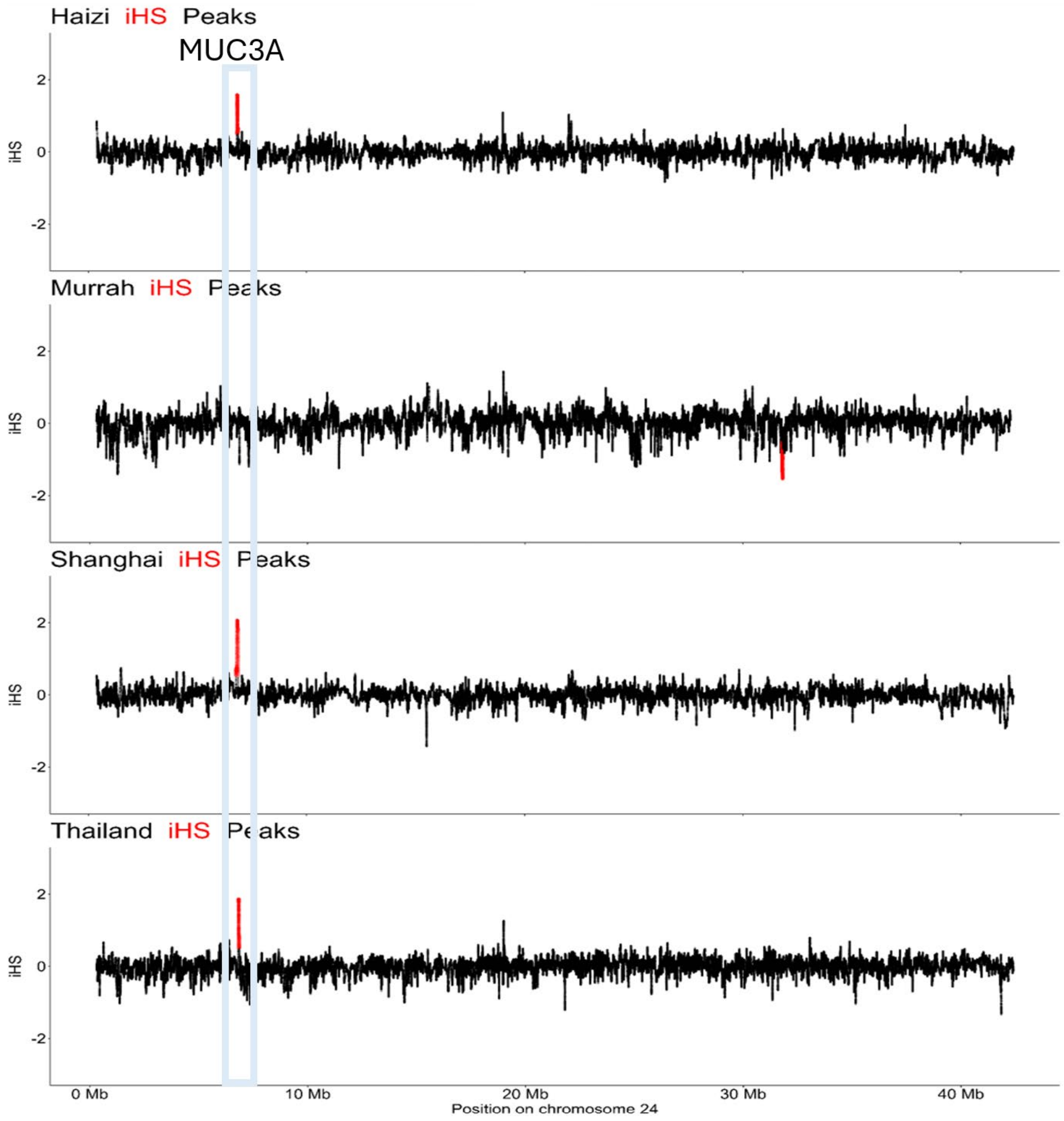


**Supplementary Fig. S10**
